# The role of intolerance of uncertainty in the acquisition and extinction of reward

**DOI:** 10.1101/2020.05.18.101212

**Authors:** Jayne Morriss, Nicolo Biagi, Tina B. Lonsdorf, Marta Andreatta

**Author notes:** Correspondence: Jayne Morriss, Centre for Integrative Neuroscience and Neurodynamics, School of Psychology and Clinical Language Sciences, University of Reading, Earley Gate, Whiteknights Campus, RG6 6AH Reading, United Kingdom.

## Abstract

Individuals, who score high in self-reported intolerance of uncertainty (IU), tend to find uncertainty anxiety-provoking. IU has been reliably associated with disrupted threat extinction. However, it remains unclear whether IU would be related to disrupted extinction to other arousing stimuli that are not threatening (i.e., rewarding). We addressed this question by conducting a reward associative learning task with acquisition and extinction training phases (*n* = 58). Throughout the associative learning task, we recorded valence ratings (i.e. liking), skin conductance response (SCR) (i.e. sweating), and corrugator supercilii activity (i.e. brow muscle indicative or negative and positive affect) to learned reward and neutral cues. During acquisition training with partial reward reinforcement, higher IU was associated with greater corrugator supercilii activity to neutral compared to reward cues. IU was not related to valence ratings or SCR’s during the acquisition or extinction training phases. These preliminary results suggest that IU-related deficits during extinction may be limited to situations with threat. The findings further our conceptual understanding of IU’s role in the associative learning and extinction of reward, and in relation to the processing of threat and reward more generally.

## 1. Introduction

Individuals, who score high in self-reported intolerance of uncertainty (IU), tend to find uncertainty negative (i.e. stressful, distressing) (Carleton, 2016a, 2016b). IU is a transdiagnostic risk factor, as high levels of self-reported IU are observed in a number of mental health disorders (Kesby, Maguire, Brownlow, & Grisham, 2017; McEvoy & Mahoney, 2012). Recent research has shown individuals high in IU, relative to individuals low in IU, to display heightened physiological and neural activity to uncertain threat and reward (for review see Morriss, Gell, & van Reekum, 2018; Tanovic, Gee, & Joormann, 2018).

The majority of the literature has, so far, focused on how IU is involved in the processing of uncertain threat. Research from our lab and others suggests that IU may play a critical role in associative threat learning (for review see Tanovic et al., 2018). During threat acquisition training with partial reinforcement, there is some evidence that individuals with high IU display greater startle blink to learned threat vs. safety cues (Chin, Nelson, Jackson, & Hajcak, 2016). However, a number of studies have not found relationships between IU and physiological measures during threat acquisition training with partial reinforcement (Dunsmoor, Campese, Ceceli, LeDoux, & Phelps, 2015; Lucas, Luck, & Lipp, 2018; Morriss, 2019). Notably, the relationship between IU and threat extinction is more consistent: during threat extinction, individuals with high IU have been shown to display greater skin conductance, pupil dilation and amygdala activity to learned threat vs. safety cues (Dunsmoor et al., 2015; Lucas et al., 2018; Morriss, Christakou, & Van Reekum, 2015, 2016; Morriss & van Reekum, 2019).

During threat extinction training, the conditioned stimulus (CS+: i.e. a shape) is presented without the associated unconditioned stimulus (US: i.e. shock or loud noise). This change in contingency may result in two different interpretations for the CS+ (Bouton, 2002): the CS+ may be interpreted either as an uncertain threat cue or as a safety cue. It has been suggested that the uncertainty surrounding the meaning of the CS during extinction is what maintains the conditioned response in individuals with high IU (Dunsmoor et al., 2015; Morriss, Saldarini, Chapman, Pollard, & van Reekum, 2019; Morriss & van Reekum, 2019). This explanation is in line with a current definition of IU, which postulates that individuals with high IU have central beliefs that information is missing or omitted, which in turn sustains the perception of uncertainty and associated heightened physiological arousal (Carleton, 2016b, p.31).

From the existing literature, it is evident that individuals with higher IU have difficulty extinguishing threat, even when mild (Dunsmoor et al., 2015; Lucas et al., 2018; Morriss, Saldarini, & Van Reekum, 2019). However, gaps still remain in the literature as to what experimental parameters induce disrupted extinction in individuals with high IU. For example, is disrupted extinction in individuals with higher IU limited to situations with uncertain threat or can it occur with other uncertain stimuli with arousing properties (i.e. rewarding)? Indeed, there is a small emerging literature on the role of IU in the processing of uncertain reward. During the anticipation of uncertain reward, individuals with high IU display greater anterior insula activity (Gorka, Nelson, Phan, & Shankman, 2016) and altered electroencephalogram and event-related potential responses (Nelson, Kessel, Jackson, & Hajcak, 2016; Nelson, Shankman, & Proudfit, 2014). Furthermore, a handful of decision-making studies have demonstrated that individuals with high IU are more likely to choose immediate smaller rewards over waiting for larger uncertain rewards (Carleton et al., 2016; Luhmann, Ishida, & Hajcak, 2011; Tanovic, Hajcak, & Joormann, 2018). Based on the aforementioned evidence, it is possible that individuals with high IU may also display differences during the acquisition and extinction of reward.

Given that IU is transdiagnostic and that associative learning principles underpin current exposure therapies (Craske, Treanor, Conway, Zbozinek, & Vervliet, 2014), examining whether IU is related to the acquisition and extinction of reward may provide crucial information relevant to the pathology and treatment of a number of mental health disorders. In particular, the findings will be relevant for work examining the effectiveness of counterconditioning (i.e. replacing threatening outcomes with rewarding outcomes) versus extinction as a potential avenue of treatment for anxiety disorders (Keller, Hennings, & Dunsmoor, 2020; Pittig, 2019). Not only will this research be useful for understanding IU in relation to anxiety disorders, but also in relation to other disorders that have a reward component and rely on exposure therapy such as substance abuse and eating disorders (Koskina, Campbell, & Schmidt, 2013; Marissen, Franken, Blanken, van den Brink, & Hendriks, 2007).

Here, we conducted an associative learning experiment to assess the relationship between individual differences in self-reported IU and conditioned reward responses. We measured skin conductance response (i.e. sweating indicative of arousal), corrugator supercilii activity (i.e. brow muscle indicative of negative or positive affect) and valence ratings (i.e. liking) throughout the acquisition and extinction training phases. We used secondary rewards (i.e., money) as unconditioned stimuli and visual shape stimuli as conditioned stimuli, in line with previous conditioning research (Andreatta & Pauli, 2015, 2019; Ebrahimi et al., 2019; Kruse, León, Stalder, Stark, & Klucken, 2018; Kruse, Tapia León, Stark, & Klucken, 2017; van den Akker, Nederkoorn, & Jansen, 2017). We used a 50% reinforcement rate during acquisition training to prolong conditioning during extinction (Bouton, 2002) and induce greater uncertainty during extinction similar to our previous work (Morriss, 2019).

We hypothesised that during reward acquisition training, skin conductance responding and valence ratings would be higher, and corrugator supercilii activity would be lower, to the learned reward (CS+) versus neutral (CS−) cues, indicative of conditioned responding (Andreatta & Pauli, 2015; Ebrahimi et al., 2019; Kruse et al., 2020; Kruse et al., 2017; van den Akker, Nederkoorn, & Jansen, 2017; Wardle, Lopez-Gamundi, & Flagel, 2018). Given the lack of IU-related findings for the acquisition training phase in past research, we did not have any specific IU hypotheses for the acquisition training phase. We hypothesised that if IU-related effects during extinction training are driven by the arousingness of an uncertain stimulus, then higher IU, relative to lower IU, will be associated with larger conditioned responding to CS+ versus CS− cues during reward extinction training. We tested the specificity of IU effects by controlling for trait anxiety, assessed by the State-Trait Inventory for Cognitive and Somatic Anxiety (STICSA) (Ree, French, MacLeod, & Locke, 2008). We selected the STICSA because it is thought to be a purer measure of anxiety, compared to other trait anxiety measures which also feature depressive symptomology (Grös, Antony, Simms, & McCabe, 2007).

## 2. Method

### 2.1 Participants

Fifty-eight participants were recruited from the University of Reading and local area through the use of advertisements and word of mouth (Sex: 29 female, 29 male; *M*_age_ = 25.42 years, *SD*_age_ = 4.70 years, range = 18-35 years; Ethnicity: 32 White, 16 Asian, 3 Mixed, 2 Black, 1 Arab, and 4 not specified; Sexual Orientation: 45 Heterosexual, 3 Sexual Minorites (homosexual/bisexual), 10 not specified). We recruited participants who were between the ages of 18-35 years. No other exclusion criteria was used for recruitment. Participants were paid £5 renumeration for their time. The procedure was reviewed and accepted by the University of Reading Research Ethics Committee.

Multilevel models (MLM) were used to analyse experimental data, where IU and STICSA scores were entered into the analysis as continuous predictor variables. Due to the complexity of MLMs, there is no agreed upon method for calculating power and estimating sample size for MLM (Peugh, 2010). Therefore, appropriate sample sizes were estimated based upon power analyses using a repeated measures ANCOVA. The sample size of this study was based on a power analysis using the average effect size (☐^2^p = .16) taken from Stimulus x Time x IU interactions for SCR magnitude from five previous experiments (4/5 with significant effects of IU)(Morriss, Christakou, & Van Reekum, 2015, 2016; Morriss & van Reekum, 2019). The following parameters were used: effect size *f* = 0.40 (converted from ☐^2^p = 0.16 and rounded down), α error probability = 0.05, Power (1-β error probability) = 0.8, numerator *df* = 1, number of covariates = 2 (IU, STICSA). The total sample size suggested was an *n* = 52. Due to non-responding in SCR (typically 5-10% of sample), we attempted to increase statistical power by recruiting a total *n* of However, due to Covid-19 recruitment was interrupted, leaving the current study with a total sample size of *n* = 58.

### 2.2 Procedure

On the day of the experiment, participants were informed about the experimental procedures upon arrival at the laboratory. Participants were then seated in the testing booth and asked to complete a consent form and a series of questionnaires (see section 2.4 below) on the computer screen.

After the questionnaires, physiological sensors were attached to the participants left hand and left corrugator supercilii. Before the task began, participants were played the sound stimulus through the headphones, so they knew what to expect. Participants were instructed: (1) to maintain attention to the task by looking at the geometrical shapes, (2) to use the keyboard for the ratings, (3) that the ‘£’ symbol and sound represented a value of £1, and (4) that the £5 from taking part was separate from the money acquired during the task, and that they would receive the total amount of money at the end of the experiment. The conditioning task (see section 2.3 below for details) was presented on a computer screen whilst skin conductance response, corrugator supercilii and valence ratings were recorded.

To maintain uncertainty, participants were not instructed about the CS-US contingency or the total amount of money that could be acquired during the task. The total amount of money that could be won during the task was fixed at £5. Therefore, all participants received a total £10 (£5 from taking part and £5 acquired from the task) at the end of the experiment.

### 2.3 Conditioning task

The conditioning task was designed using E-Prime 2.0 software (Psychology Software Tools Ltd, Pittsburgh, PA). Participants were sat approximately 60 cm from the screen. Visual stimuli were yellow and blue squares displayed on a black background. To represent monetary reward, the presentation of a ‘£’ symbol and a 1000 ms 70 dB sound of coins dropping served as the US (Kruse, Klein, Tapia León, Stark, & Klucken, 2020; Kruse et al., 2018; Kruse et al., 2017; Tapia León, Kruse, Stark, & Klucken, 2019).

The task comprised of two phases: acquisition and an immediate extinction training. During acquisition training, one of the squares was paired with the US 50% of the time (CS+), whilst the other square was presented alone (CS−). During extinction training, the squares were presented in the absence of the US. There was no break between the acquisition and extinction training phases. Conditioning contingencies were counterbalanced.

The acquisition training phase consisted of 16 trials (4 CS+ paired, 4 CS+ unpaired, 8 CS−). The extinction training phase comprised of 32 trials (16 CS+ and 16 CS−), where early is defined as the first 8 CS+/CS− trials and late is defined as the last 8 CS+/CS− trials. Experimental trials were pseudo-randomised such that the first trial of acquisition training was always paired and then after each trial type could only be played 3 times in a row. The squares were presented for a total of 6 seconds. After this, a blank black screen was presented for 8-12 seconds. During reinforced trials, presentation of the square coterminated exactly with the presentation of the US.

Participants were presented with two other 9-point Likert scales at the end of the experiment. Participants were asked to rate: (1) the valence and (2) the arousal of the stimuli (i.e. the ‘£’ symbol and sound stimulus combined). These scales ranged from 1 (Valence: very negative; Arousal: calm) to 9 (Valence: very positive; Arousal: excited).

### 2.4 Questionnaires

To assess intolerance of uncertainty and trait anxiety, we administered the Intolerance of Uncertainty Scale (IUS) (Freeston, Rhéaume, Letarte, Dugas, & Ladouceur, 1994) and State-Trait Inventory for Cognitive and Somatic Anxiety (STICSA) (Ree, French, MacLeod, & Locke, 2008). The IUS measure consists of 27 items that are rated on a 5-point Likert scale. The STICSA measure consists of 21 items that are rated on a 4-point Likert scale.

### 2.5 Skin conductance acquisition and scoring

Identical to previous work (Morriss, 2019), physiological recordings were obtained using AD Instruments (AD Instruments Ltd, Chalgrove, Oxfordshire) hardware and software. Electrodermal activity was measured with dry MLT116F silver/silver chloride bipolar finger electrodes that were attached to the distal phalanges of the index and middle fingers of the left hand. A low constant-voltage AC excitation of 22 mVrms at 75 Hz was passed through the electrodes, which was connected to a ML116 GSR Amp, and converted to DC before being digitized and stored. A PowerLab 26T Unit Model was used to amplify the skin conductance signal, which was digitized through a 16-bit A/D converter at 1000 Hz. The electrodermal signal was converted from volts to microSiemens using AD Instruments software (AD Instruments Ltd, Chalgrove, Oxfordshire).

Skin conductance response onsets and offsets were marked using ADinstruments software (AD Instruments Ltd, Chalgrove, Oxfordshire) and extracted using Matlab R2017a software (The MathWorks, Inc., Natick, Massachusetts, United States). Skin conductance response onsets and offsets were assigned using a macro in ADinstruments and then visually inspected to ensure the onsets and offsets were assigned correctly. Skin conductance responses (SCR) were scored when there was an increase of skin conductance level exceeding 0.03 microSiemens (Dawson, Schell, & Filion, 2000). The amplitude of each response was scored as the difference between the onset and the maximum deflection prior to the signal flattening out or decreasing. SCR onsets and respective peaks were counted if the SCR onset was within 0.5-4 seconds (CS response) following CS onset (Bauer et al., 2020; Morriss, Macdonald, & van Reekum, 2016). Trials with no discernible SCRs were scored as zero. SCR magnitudes were then square root transformed to reduce skew and *z*-scored to control for interindividual differences in skin conductance responsiveness (Ben‐Shakhar, 1985). SCR magnitudes were calculated from remaining trials by averaging SCR-transformed values for each condition (Acquisition CS+; Acquisition CS−; Extinction Learning CS+ Early; Extinction Learning CS− Early; Extinction Learning CS+ Late; Extinction Learning CS− Late). Non-responders were defined as those who responded to 10% or less of the total CS+ and CS− trials across acquisition and extinction training (48 trials in total) (Morriss, Chapman, Tomlinson, & Van Reekum, 2018; Xia, Dymond, Lloyd, & Vervliet, 2017). Using this criterion, 2 non-responders were excluded from the SCR analyses, leaving 56 participants with useable SCR data.

### 2.6 Corrugator supercilii acquisition and scoring

The protocol for corrugator acquisition and scoring was in line with previous research (Morriss, 2019). Facial electromyography measurement of the left corrugator supercilii was obtained by using a pair of 4 mm Ag/AgCl bipolar surface electrodes connected to the ML138 Bio Amp. The bipolar surface electrodes were placed approximately 15 mm apart. The reference electrode was a singular 8 mm Ag/AgCl electrode, placed upon the middle of the forehead, and connected to the ML138 Bio Amp. Before placing the sensors, the skin site was slightly abraded with isopropyl alcohol skin prep pads to reduce skin impedance to an acceptable level (below 20 kΩ). EMG was sampled at 1000 Hz. A high‐pass filter of 20 Hz was applied to the raw EMG online (Solnik, DeVita, Rider, Long, & Hortobágyi, 2008). The EMG were root mean squared offline (Fridlund & Cacioppo, 1986).

Corrugator supercilii activity was extracted using R software (R Core Team, 2014). Corrugator supercilii activity was averaged for each 1000 ms window following CS onset, resulting in six windows of 1000 ms each. These data were baseline corrected by subtracting 2000 ms preceding each CS onset from a blank screen (similar to procedures used by Morriss, 2019). Corrugator supercilii trials were averaged per condition and second window (Acquisition CS+; Acquisition CS−; Extinction Learning CS+ Early; Extinction Learning CS− Early; Extinction Learning CS+ Late; Extinction Learning CS− Late).

For the corrugator supercilii data, there was a recording error for 1 participant, thus, leaving 57 participants with useable corrugator supercilii data.

### 2.7 Analyses

The analyses were conducted using the mixed procedure in SPSS 24.0 (SPSS, Inc; Chicago, Illinois). We conducted separate MLMs for SCR magnitude, corrugator supercilii activity and valence ratings during acquisition and extinction training. For SCR magnitude and valence ratings during the acquisition training phase we entered Stimulus (CS+, CS−) at level 1 and individual subjects at level 2. For SCR magnitude and valence ratings during the extinction training phase we entered Stimulus (CS+, CS−) and Time (Early: first 8 CS+/CS− trials, Late: last 8 CS+/CS− trials) at level 1 and individual subjects at level 2. For corrugator supercilii activity, an additional factor of Second (time bins: 1,2,3,4,5,6) at level 1 was included in the MLMs. We included individual difference predictor variables (IU and STICSA) in the MLMs. In all models, we used a diagonal covariance matrix for level 1. Random effects included a random intercept for each individual subject, where a variance components covariance structure was used. Fixed effects included Stimulus and Time. We used a maximum likelihood estimator for the MLMs.

The IUS and STICSA covariates were entered separately. If there was a significant interaction with one of the predictor variables (IUS, STICSA), then we conducted a further MLM with both predictor variables entered to test specificity. Based on previous work (Morriss, 2019), we expected such specificity for IUS, but we explored interactions with STICSA, given findings with trait anxiety in the conditioning literature (Lonsdorf & Merz, 2017). Where a significant interaction was observed with IUS (or STICSA), we performed follow-up pairwise comparisons on the estimated marginal means of the relevant conditions estimated at specific IUS values of + or −1 SD of mean IUS, adjusted for STICSA (or IUS) (Morriss, 2019; Morriss, Saldarini, & Van Reekum, 2019).

## 3. Results

For descriptive statistics see Table 1.

**Table 1.** Summary of means (SD) for each dependent measure as a function of condition (CS+ and CS−), separately for acquisition, early extinction and late extinction.

| Measure | Acquisition |  | Early Extinction |  | Late Extinction |  |
| --- | --- | --- | --- | --- | --- | --- |
|  | CS+ | CS- | CS+ | CS- | CS+ | CS- |
| Square root transformed and z-scored SCR magnitude ($\sqrt{\mu\text{S}}$) | .05 (-.38) | -.07 (.31) | .04 (.40) | -.12 (.27) | .01 (.34) | .02 (.40) |
| Z-scored corrugator supercilii activity ($\mu\text{V}$) | -.01 (-.23) | .01 (.23) | -.01 (0.22) | -.01 (0.26) | .04 (.25) | -.05 (.25) |
| Liking rating (1-9) | 6.34 (1.93) | 3.84 (2.11) | 4.86 (1.96) | 3.64 (2.19) | 3.71 (1.91) | 3.33 (2.08) |

Note: SCR magnitude ( $\sqrt{\mu\text{S}}$ ), square root transformed and z-scored skin conductance magnitude measured in microSiemens. Corrugator supercilii activity ( $\mu\text{V}$ ), z-scored corrugator supercilii activity measured in microVolts. Liking ratings from 1-9, where 1 is dislike and 9 is like.

### 3.1 Questionnaires

The self-reported anxiety measures had excellent internal reliability: IUS (*M* = 63.87, *SD* = 20.06, range= 30-112, *α* = .94); STICSA (*M* = 38.43, *SD* = 9.88, range = 24-66, *α* = .89). IUS was positively significantly correlated with STICSA, *r*(56) = .745, *p* < .001.

### 3.2 Ratings

The ‘£’ symbol and sound stimulus was on average rated as moderately positive (*M* = 7.39 *SD* = 1.58, range 3-9, where 1 = very negative and 9 = very positive) and moderately arousing (*M* = 6.36, *SD* = 2.10, 1-9, range where 1 = calm and 9 = excited).

During acquisition training participants reported liking the CS+, compared to the CS− [Stimulus: *F*(1, 115.186) = 47.372, *p* < .001; see Table 1 and Figure 1]. No significant interactions with IUS or STICSA were observed for the ratings during acquisition training, max *F* =1.332.

**Figure 1.**
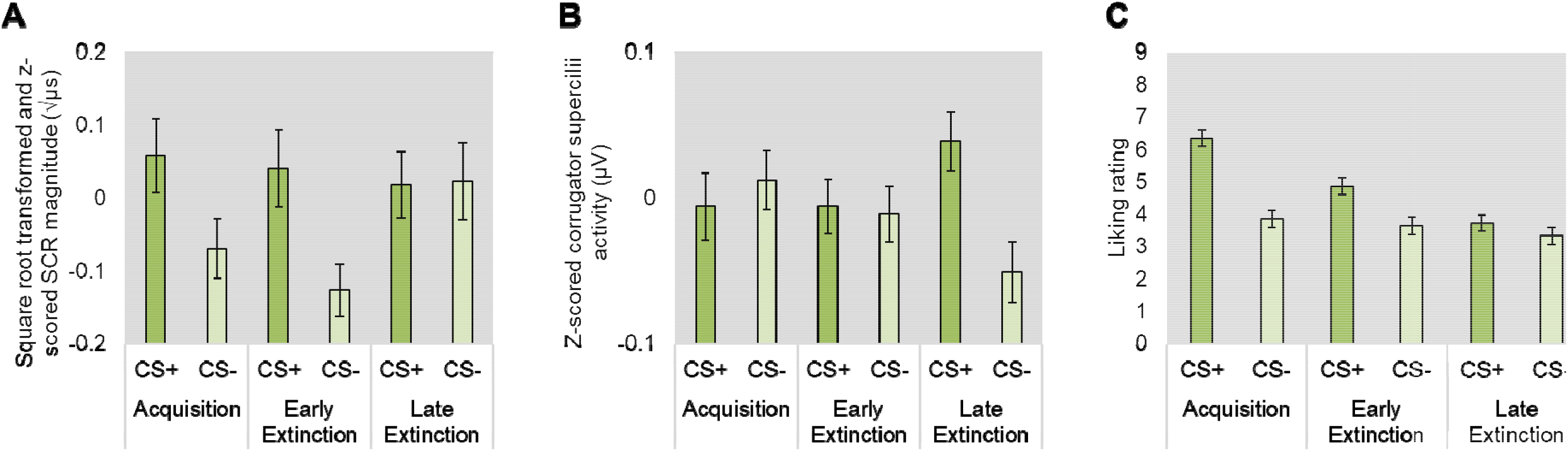
Bar graphs depicting mean (A) SCR magnitude, (B) corrugator supercilii activity, and (C) liking ratings for each stimulus type during each experimental phase of the associative reward learning task. Error bars represent standard error. Expectancy ratings, 1 = dislike, 9 = like. Square root transformed and z-transformed SCR magnitude (μS), skin conductance magnitude measured in microSiemens. Z-scored corrugator supercilii activity (μV), measured in microVolts.

During extinction training, participants reported: (1) neither liking or disliking the CS+, and (2) disliking the CS− [Stimulus: *F*(1, 170.616) = 16.327, *p* < .001; see Table 1 and Figure 1]. Ratings of liking dropped over time [Time: *F*(1, 170.616) = 13.639, *p* = .*p* < .001; Stimulus x Time: *F*(1, 170.616) = 4.533, *p* = .035]. Follow-up pairwise comparisons revealed that participants reported: (1) liking the CS+ more than the CS− during early extinction training, *p* < .001, but not during late extinction training, *p* = .165; (2) liking the CS+ more during early, compared to late extinction training, *p* < .001; and (3) disliking the CS− similarly during early and late extinction training, *p* = .295. No significant interactions with IUS or STICSA were found for the ratings during extinction training, max *F* = 2.755.

### 3.3 SCR

As expected, during acquisition training SCR’s were larger to the CS+, compared to the CS− [Stimulus: *F*(1, 56) = 4.099, *p* = .048; see Table 1 and Figure 1]. No significant interactions with IUS or STICSA were observed for SCR’s during acquisition training, max *F* = 2.606.

During extinction training, there were no significant effects of Stimulus, Time or Stimulus x Time, or interactions with IUS and STICSA, max *F* = 3.406. Although, SCR was larger to the CS+ vs. CS− during early extinction training at trend level [Stimulus x Time: *F*(1,207.949) = 3.406, *p* = .066; see Table 1 and Figure 1].

### 3.4 Corrugator supercilii activity

During acquisition training, we did not observe a significant difference for corrugator supercilii activity to the CS+, versus the CS− [Stimulus: *F*(1, 471.238) = .970, *p* = .325; see Table 1]. However, during acquisition training, IUS scores were associated with differences in corrugator supercilii activity to the CS+ and CS− [Stimulus x IUS: *F*(1, 477.218) = 12.065, *p* = .001]. Importantly, the interaction remained significant when STICSA was entered into the MLM [Stimulus x IUS: *F*(1, 471.238) = 6.126, *p* = .014; see Figure 2]. Follow up pairwise comparisons revealed that individuals with higher IUS scores showed greater corrugator supercilii activity to the CS−, compared to the CS+, *p* = .011, while individuals with lower IUS scores did not show differences in corrugator supercilii activity to the CS+, compared to the CS−, *p* =.113. Further post hoc correlations between IUS and the CS+ and CS− separately confirmed that higher IUS was associated with greater corrugator supercilii activity to the CS− [*r*(55) = .257, *p* = .053] but not the CS+ [*r*(55) = −.074, *p* = .583]^1^.

**Figure 2.**
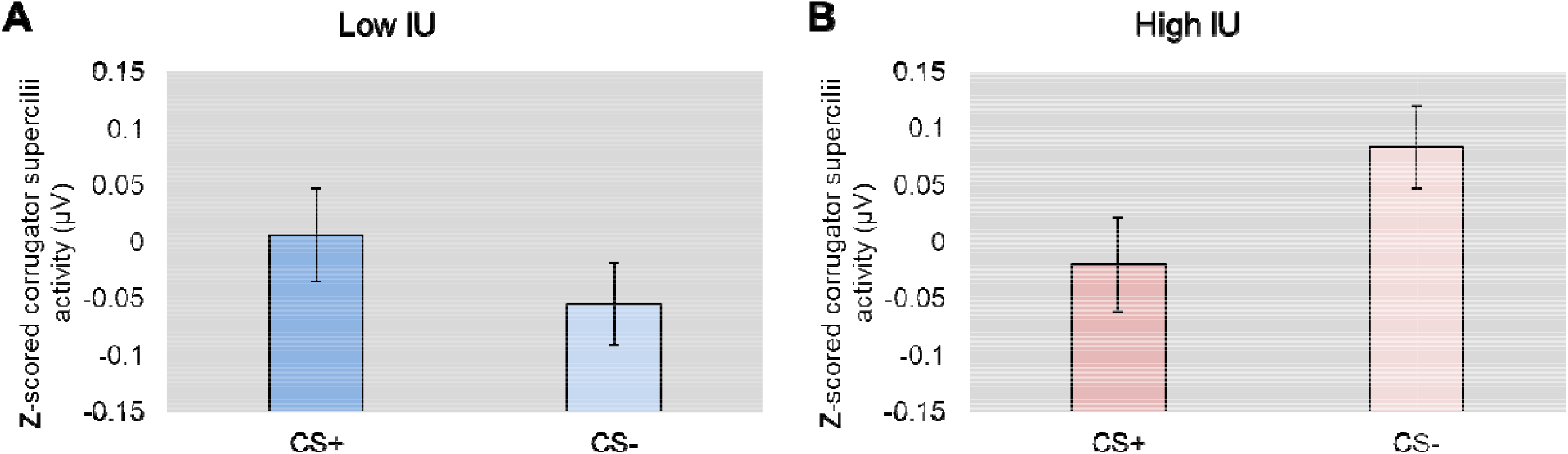
Bar graphs depicting IU estimated at + or − 1 SD of mean IU (controlling for STICSA) from the multilevel model analysis for corrugator supercilii activity during the acquisition phase of the associative reward learning task (A and B). Higher IU, relative to lower IU, was associated with greater corrugator supercilii activity to the CS−, compared to the CS+ during reward acquisition. Bars represent standard error at + or – 1 SD of mean IU. Z-scored corrugator supercilii activity (μV), measured in microVolts.

We observed no differences in corrugator supercilii activity to the CS+, versus the CS− during early extinction training, *p* = .831. However, corrugator supercilii activity was greater to the CS+ versus CS− during late extinction training, *p* = .001 [Stimulus: *F*(1, 1198.488) = 6.511, *p* = .011; Stimulus x Time: *F*(1, 1198.488) = 4.892, *p* = .027; see Table 1 and Figure 1]. No significant interactions with IUS or STICSA were observed for the corrugator supercilii during extinction training, max *F* = 1.192.

## 4. Discussion

In the current study, we examined the role of self-reported IU in reward acquisition and extinction. IU was related to greater corrugator supercilii activity to neutral vs. reward cues during acquisition training. However, IU was not related to any other measure during the reward acquisition or extinction training phases. These results further our understanding of the role of IU in the associative learning of reward, beyond the domain of threat.

In line with previous work, reward acquisition was observed via larger SCR magnitude and liking ratings to the learned reward vs. neutral cues (Andreatta & Pauli, 2015, 2019; Ebrahimi et al., 2019; Kruse et al., 2018; Kruse et al., 2017; Tapia León et al., 2019; Wardle et al., 2018). However, reward acquisition was not observed via the corrugator supercilii. We would have expected less corrugator supercilii activity to the reward vs. neutral cues, indicative of positive affect towards the reward cues (Wardle et al., 2018). Furthermore, typical patterns of reward extinction were observed: larger SCR magnitude and liking ratings were found for the early part of the extinction training phase, relative to the late part of extinction training phase (Andreatta & Pauli, 2015; Ebrahimi et al., 2019; Kruse et al., 2020; Kruse et al., 2017). Interestingly, during late extinction training there was larger corrugator supercilii to the reward cues compared to the neutral cues. We speculate that this corrugator supercilii result may reflect greater negative affect to cues previously associated with reward during extinction training, due to the omission (or loss) of reward.

Unexpectedly, during reward acquisition training, we found that higher IU, over STICSA, was specifically related to greater corrugator supercilii activity to neutral vs. reward cues. Moreover, lower IU was associated with less corrugator supercilii to both neutral and reward cues. Given that the corrugator supercilii is thought to reflect negative affect (i.e. fear, anxiety, distress, irritation, pain), positive affect and even effort (Larsen, Norris, & Cacioppo, 2003; Tassinary, Cacioppo, & Vanman, 2007), the IU-related results reported here may have multiple interpretations. Firstly, the corrugator supercilii results may reflect differences in effort: individuals high in IU, relative to low IU may have been more motivated (i.e. effort) to keep track of the contingencies because the reward was unpredictable. Secondly, the corrugator supercilii results may reflect differences in valence: individuals high IU, relative to low IU, may have found the neutral vs. reward cues more negative because they weren’t associated with a rewarding outcome. While, we did not hypothesise the IU-related effect during reward acquisition training, it is in agreement with the broader literature on IU and uncertain reward (Carleton et al., 2016; Gorka et al., 2016; Luhmann et al., 2011; Tanovic, Hajcak, et al., 2018). Further work is needed to assess the reliability of this IU-related result during reward acquisition training. For example, does the effect vary depending on the level of uncertainty (i.e. reinforcement rate) and the type of reward (i.e. money, food, substances)?

The hypothesis was that if IU-related effects during extinction are driven by the arousingness of an uncertain stimulus, then higher IU, relative to lower IU, would be associated with larger conditioned responding to reward vs. neutral cues during extinction training. However, IU was not associated with any of the rating or physiological measures during reward extinction training. In relation to the broader literature on IU and threat extinction, the results from the current experiment suggest that IU-related deficits in extinction are limited to situations with uncertain threat (Dunsmoor et al., 2015; Morriss, 2019; Morriss & van Reekum, 2019) and may not occur in the same way for situations with uncertain reward, despite potential similarity in arousingness. Importantly, however, this is only a single study and further evidence is warranted to make stronger conclusions on the role of IU in reward extinction. For example, perhaps, we would observe IU-related effects during reward extinction if the reward was more arousing (i.e. larger monetary rewards) or motivationally relevant in relation to a threat or loss to the self (i.e. money when poorer, food when hungry, and substances such as nicotine or alcohol when addicted).

In the current study we show that individuals with high IU are able to acquire and extinguish uncertain reward. These results may have implications for researchers examining counterconditioning (i.e. replacing threatening outcomes with rewarding outcomes) as an alternative to threat extinction training (i.e. replacing threatening outcomes with nothing) (Keller, Hennings, & Dunsmoor, 2020; Pittig, 2019) more broadly, and in relation to IU. Counterconditioning may be more effective than standard extinction training for individuals with high IU because during counterconditioning the new reward association is explicitly reinforced (i.e. a reward is given), whereas during standard extinction training the new association is less obvious (i.e. nothing happens). On this basis, counterconditioning vs. standard extinction training may be less distressing and therefore lead to more effective removal of old threat associations in individuals with high IU.

In conclusion, higher IU was associated with greater corrugator supercilii activity to neutral vs. reward cues during acquisition training. However, IU was not related to any other measure during the reward acquisition or extinction training phases. These initial results further our conceptual understanding of IU in reward associative learning, and in relation to the processing of reward and threat more generally. In order to assess the generalisability and reliability of the results reported here, further research is needed to examine how individual differences in IU modulate reward acquisition and extinction (i.e. vary levels of uncertainty and reward).

## Acknowledgements

This research was supported by: (1) a NARSAD Young Investigator Grant from the Brain & Behavior Research Foundation (27567) and an ESRC New Investigator Grant (ES/R01145/1) awarded to Jayne Morriss. To access the data, please contact Dr. Jayne Morriss.

## Conflict of interest statement

The authors state no conflict of interest.

## Footnotes

1 Results for the corrugator supercilii during acquisition training were similar when the IU-12 scale was used (Carleton, Norton, & Asmundson, 2007) [MLM: Stimulus x IU-12: *F*(1, 467.447) = 8.658, *p* = .003; Correlations between IU-12 and CS− [*r*(55) = .297, *p* = .025], and CS+ [*r*(55) = −.065, *p* = .633]].

## Notes

### Competing Interest Statement

The authors have declared no competing interest.

## References

Andreatta, M., & Pauli, P. (2015). Appetitive vs. aversive conditioning in humans. Frontiers in Behavioral Neuroscience, 9, 128.

Andreatta, M., & Pauli, P. (2019). Generalization of appetitive conditioned responses. Psychophysiology, e13397.

Bauer, E. A., MacNamara, A., Sandre, A., Lonsdorf, T. B., Weinberg, A., Morriss, J., & Van Reekum, C. M. (2020). Intolerance of uncertainty and threat generalization: A replication and extension. Psychophysiology, e13546.

Ben‐Shakhar, G. (1985). Standardization within individuals: A simple method to neutralize individual differences in skin conductance. Psychophysiology, 22(3), 292–299.

Bouton, M. E. (2002). Context, ambiguity, and unlearning: Sources of relapse after behavioral extinction. Biological Psychiatry, 52(10), 976–986.

Carleton, R. N. (2016a). Fear of the unknown: One fear to rule them all? Journal of Anxiety Disorders, 41, 5–21.

Carleton, R. N. (2016b). Into the unknown: A review and synthesis of contemporary models involving uncertainty. Journal of Anxiety Disorders, 39, 30–43.

Carleton, R. N., Duranceau, S., Shulman, E. P., Zerff, M., Gonzales, J., & Mishra, S. (2016). Self-reported intolerance of uncertainty and behavioural decisions. Journal of Behavior Therapy and Experimental Psychiatry, 51, 58–65.

Carleton, R. N., Norton, M. P. J., & Asmundson, G. J. (2007). Fearing the unknown: A short version of the Intolerance of Uncertainty Scale. Journal of Anxiety Disorders, 21(1), 105–117.

Chin, B., Nelson, B. D., Jackson, F., & Hajcak, G. (2016). Intolerance of uncertainty and startle potentiation in relation to different threat reinforcement rates. International Journal of Psychophysiology, 99, 79–84.

Craske, M. G., Treanor, M., Conway, C. C., Zbozinek, T., & Vervliet, B. (2014). Maximizing exposure therapy: an inhibitory learning approach. Behaviour Research and Therapy, 58, 10–23.

Dawson, M. E., Schell, A. M., & Filion, D. L. (2000). The Electrodermal System. In J. T. Cacioppo, L. G. Tassinary, & G. G. Berntson (Eds.), Handbook of Physiology (2nd ed., pp. 200–223). Cambridge, UK: Cambridge University Press.

Dunsmoor, J. E., Campese, V. D., Ceceli, A. O., LeDoux, J. E., & Phelps, E. A. (2015). Novelty-facilitated extinction: providing a novel outcome in place of an expected threat diminishes recovery of defensive responses. Biological Psychiatry, 78(3), 203–209.

Ebrahimi, C., Koch, S. P., Pietrock, C., Fydrich, T., Heinz, A., & Schlagenhauf, F. (2019). Opposing roles for amygdala and vmPFC in the return of appetitive conditioned responses in humans. Translational Psychiatry, 9(1), 1–12.

Freeston, M. H., Rhéaume, J., Letarte, H., Dugas, M. J., & Ladouceur, R. (1994). Why do people worry? Personality and Individual Differences, 17(6), 791–802.

Fridlund, A. J., & Cacioppo, J. T. (1986). Guidelines for human electromyographic research. Psychophysiology, 23(5), 567–589.

Gorka, S. M., Nelson, B. D., Phan, K. L., & Shankman, S. A. (2016). Intolerance of uncertainty and insula activation during uncertain reward. Cognitive, Affective, & Behavioral Neuroscience, 16(5), 929–939.

Grös, D. F., Antony, M. M., Simms, L. J., & McCabe, R. E. (2007). Psychometric properties of the State-Trait Inventory for Cognitive and Somatic Anxiety (STICSA): comparison to the State-Trait Anxiety Inventory (STAI). Psychological Assessment, 19(4), 369.

Keller, N. E., Hennings, A. C., & Dunsmoor, J. E. (2020). Behavioral and neural processes in counterconditioning: Past and future directions. Behaviour Research and Therapy, 125, 103532.

Kesby, A., Maguire, S., Brownlow, R., & Grisham, J. R. (2017). Intolerance of uncertainty in eating disorders: an update on the field. Clinical Psychology Review, 56, 94–105.

Koskina, A., Campbell, I. C., & Schmidt, U. (2013). Exposure therapy in eating disorders revisited. Neuroscience & Biobehavioral Reviews, 37(2), 193–208.

Kruse, O., Klein, S., Tapia León, I., Stark, R., & Klucken, T. (2020). Amygdala and nucleus accumbens involvement in appetitive extinction. Human Brain Mapping, 41(7).

Kruse, O., León, I. T., Stalder, T., Stark, R., & Klucken, T. (2018). Altered reward learning and hippocampal connectivity following psychosocial stress. Neuroimage, 171, 15–25.

Kruse, O., Tapia León, I., Stark, R., & Klucken, T. (2017). Neural correlates of appetitive extinction in humans. Social Cognitive and Affective Neuroscience, 12(1), 106–115.

Larsen, J. T., Norris, C. J., & Cacioppo, J. T. (2003). Effects of positive and negative affect on electromyographic activity over zygomaticus major and corrugator supercilii. Psychophysiology, 40(5), 776–785.

Lonsdorf, T. B., & Merz, C. J. (2017). More than just noise: Inter-individual differences in fear acquisition, extinction and return of fear in humans-Biological, experiential, temperamental factors, and methodological pitfalls. Neuroscience & Biobehavioral Reviews, 80, 703–728.

Lucas, K., Luck, C. C., & Lipp, O. V. (2018). Novelty-facilitated extinction and the reinstatement of conditional human fear. Behaviour Research and Therapy, 109, 68–74.

Luhmann, C. C., Ishida, K., & Hajcak, G. (2011). Intolerance of uncertainty and decisions about delayed, probabilistic rewards. Behavior Therapy, 42(3), 378–386.

Marissen, M. A., Franken, I. H., Blanken, P., van den Brink, W., & Hendriks, V. M. (2007). Cue exposure therapy for the treatment of opiate addiction: results of a randomized controlled clinical trial. Psychotherapy and Psychosomatics, 76(2), 97–105.

McEvoy, P. M., & Mahoney, A. E. (2012). To be sure, to be sure: Intolerance of uncertainty mediates symptoms of various anxiety disorders and depression. Behavior Therapy, 43(3), 533–545.

Morriss, J. (2019). What do I do now? Intolerance of uncertainty is associated with discrete patterns of anticipatory physiological responding to different contexts. Psychophysiology, e13396.

Morriss, J., Chapman, C., Tomlinson, S., & Van Reekum, C. M. (2018). Escape the bear and fall to the lion: The impact of avoidance availability on threat acquisition and extinction. Biological Psychology, 138, 73–80.

Morriss, J., Christakou, A., & Van Reekum, C. M. (2015). Intolerance of uncertainty predicts fear extinction in amygdala-ventromedial prefrontal cortical circuitry. Biology of Mood & Anxiety Disorders, 5(1), 1.

Morriss, J., Christakou, A., & Van Reekum, C. M. (2016). Nothing is safe: Intolerance of uncertainty is associated with compromised fear extinction learning. Biological Psychology, 121, 187–193.

Morriss, J., Gell, M., & van Reekum, C. M. (2018). The uncertain brain: A co-ordinate based meta-analysis of the neural signatures supporting uncertainty during different contexts. Neuroscience & Biobehavioral Reviews, 96, 241–249.

Morriss, J., Macdonald, B., & van Reekum, C. M. (2016). What Is Going On Around Here? Intolerance of Uncertainty Predicts Threat Generalization. PloS one, 11(5), e0154494.

Morriss, J., Saldarini, F., Chapman, C., Pollard, M., & van Reekum, C. M. (2019). Out with the old and in with the new: The role of intolerance of uncertainty in reversal of threat and safety. Journal of Experimental Psychopathology, 10(1), 2043808719834451.

Morriss, J., Saldarini, F., & Van Reekum, C. M. (2019). The role of threat level and intolerance of uncertainty in extinction. International Journal of Psychophysiology, 142, 1–9.

Morriss, J., & van Reekum, C. M. (2019). I feel safe when i know: Contingency instruction promotes threat extinction in high intolerance of uncertainty individuals. Behaviour Research and Therapy, 116, 111–118.

Nelson, B. D., Kessel, E. M., Jackson, F., & Hajcak, G. (2016). The impact of an unpredictable context and intolerance of uncertainty on the electrocortical response to monetary gains and losses. Cognitive, Affective, & Behavioral Neuroscience, 16(1), 153–163.

Nelson, B. D., Shankman, S. A., & Proudfit, G. H. (2014). Intolerance of uncertainty mediates reduced reward anticipation in major depressive disorder. Journal of Affective Disorders, 158, 108–113.

Peugh, J. L. (2010). A practical guide to multilevel modeling. Journal of School Psychology, 48(1), 85–112.

Pittig, A. (2019). Incentive-based extinction of safety behaviors: Positive outcomes competing with aversive outcomes trigger fear-opposite action to prevent protection from fear extinction. Behaviour Research and Therapy, 121, 103463.

Ree, M. J., French, D., MacLeod, C., & Locke, V. (2008). Distinguishing cognitive and somatic dimensions of state and trait anxiety: Development and validation of the State-Trait Inventory for Cognitive and Somatic Anxiety (STICSA). Behavioural and Cognitive Psychotherapy, 36(3), 313–332.

Solnik, S., DeVita, P., Rider, P., Long, B., & Hortobágyi, T. (2008). Teager–Kaiser Operator improves the accuracy of EMG onset detection independent of signal-to-noise ratio. Acta of bioengineering and biomechanics/Wroclaw University of Technology, 10(2), 65.

Tanovic, E., Gee, D. G., & Joormann, J. (2018). Intolerance of uncertainty: Neural and psychophysiological correlates of the perception of uncertainty as threatening. Clinical Psychology Review, 60, 87–99.

Tanovic, E., Hajcak, G., & Joormann, J. (2018). Hating waiting: Individual differences in willingness to wait in uncertainty. Journal of Experimental Psychopathology, 9(1), 2043808718778982.

Tapia León, I., Kruse, O., Stark, R., & Klucken, T. (2019). Relationship of sensation seeking with the neural correlates of appetitive conditioning. Social Cognitive and Affective Neuroscience, 14(7), 769–775.

Tassinary, L. G., Cacioppo, J. T., & Vanman, E. J. (2007). The skeletomotor system: Surface electromyography.

van den Akker, K., Nederkoorn, C., & Jansen, A. (2017). Electrodermal responses during appetitive conditioning are sensitive to contingency instruction ambiguity. International Journal of Psychophysiology, 118, 40–47.

Wardle, M. C., Lopez-Gamundi, P., & Flagel, S. B. (2018). Measuring appetitive conditioned responses in humans. Physiology & Behavior, 188, 140–150.

Xia, W., Dymond, S., Lloyd, K., & Vervliet, B. (2017). Partial reinforcement of avoidance and resistance to extinction in humans. Behaviour Research and Therapy, 96, 79–89.

